# Per-pixel unmixing of spectrally overlapping fluorophores using intra-exposure excitation modulation

**DOI:** 10.1101/2023.04.29.538742

**Authors:** Hana Valenta, Franziska Bierbuesse, Raffaele Vitale, Cyril Ruckebusch, Wim Vandenberg, Peter Dedecker

## Abstract

Multilabel fluorescence imaging is essential for the visualization of complex systems, though a major challenge is the limited width of the usable spectral window. Here, we present a new method, exNEEMO, that enables per-pixel quantification of spectrally-overlapping fluorophores based on their light-induced dynamics, in a way that is compatible with a very broad range of timescales over which these dynamics may occur. Our approach makes use of intra-exposure modulation of the excitation light to distinguish the different emitters given their reference responses to this modulation. We use approach to simultaneously image four green photochromic fluorescent proteins at the full spatial resolution of the imaging.

**Graphical abstract:** 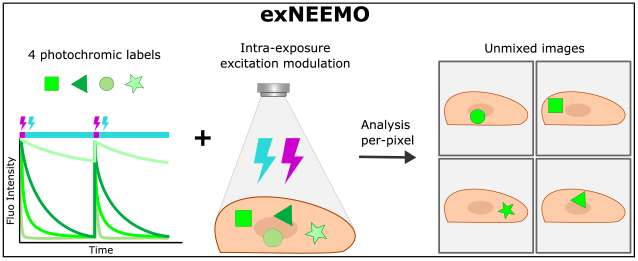

## Introduction

Fluorescence microscopy is a key technology in life sciences, though it is limited in the number of components that can be visualized at once. Several solutions to address this limitation have been developed, including computational approaches based on spectral unmixing [1] or the use of fluorescence properties other than the emission color, such as fluorescence anisotropy [2], fluorescence lifetime [3], or light-induced processes resulting in characteristic fluorescence dynamics in the emission [4, 5, 6, 7].

We recently developed emitter separation based on intra-exposure excitation modulation [8], a method that can distinguish multiple spectrally-identical fluorophores based on differences in their light-induced fluorescence dynamics that can occur over a broad range of timescales. Using this approach, which we retroactively named intra-exposure excitation modulation (NEEMO), we were able to distinguish the light-induced dynamics of four spectrally-identical reversibly photoswitchable fluorescent proteins (rsFPs) expressed within *E. coli* bacteria. Such labels can be controllably switched between a fluorescent and a non-fluorescent state using illumination at two different wavelengths.

This previous work focused on the classification of cells expressing just one of these fluorophores rather than on the separation of the fluorophore contributions within potentially complex mixtures. Classifications are easier to achieve since the signal can be averaged over the entire cell geometry, leading to much better signal-to-noise ratios, while the analysis can be based on clustering approaches that are tolerant to changes in label behavior as long as each cell can still be assigned to the correct cluster. Such an approach has also been used to, for example, distinguish up to 9 among 16 spectrally similar fluorophores expressed in bacteria [9].

The true quantitative separation of fluorophores based on their fluorescence dynamics has also seen multiple efforts over the past years, resulting in approaches such as OLID, SAFIRe, OPIOM, and HIGHLIGHT [10, 11, 12, 13, 9, 14], that make use of the direct detection of the fluorescence dynamics by the microscope. This may, however, be challenging if the detection is done using a camera that is typically much slower than the point detectors used in confocal imaging, meaning that only correspondingly slow fluorescence dynamics can be used.

In this manuscript, we explored whether our previous concept can be extended to provide a quantitative emitter separation in complex mixtures, in a manner that takes advantage of the full spatial resolution of the imaging while still being capable of leveraging fluorescence dynamics that occur too quickly to be captured by the detector. In particular, we present the extended intra-exposure excitation modulation (exNEEMO) method that permits the per-pixel separation of four spectrally-identical rsFPs based on their unique photo-switching profiles, using the acquisition of just four fluorescence images. We find that this can be successfully achieved while retaining compatibility with a broad range of light-induced fluorescence dynamics, though the overall accuracy is limited by cell-to-cell variations in these dynamics. Overall, our approach enables new dimensions for the multiplexed fluorescence imaging of complex biological systems.

## Results and discussion

We start by briefly revisiting the principle underlying the NEEMO method, schematically shown in Figure 1A and previously reported in [8]. The core idea is that different photoswitching efficiencies of the fluorophores lead to characteristic emission intensity profiles in response to temporally-varying illumination intensities. In NEEMO, we do not detect these responses directly, but instead measure only their integrated emission over the camera exposure time. The different labels can then be identified by observing how the integrated intensities vary for different illumination patterns [8]. This approach is compatible with a very broad range of processes since we do not require that the dynamics are directly resolvable by the imaging system. Instead, we require only that they can be probed within the modulation rate of the light sources, which is typically very fast (within microseconds or faster) even for inexpensive light sources.

**Figure 1.**
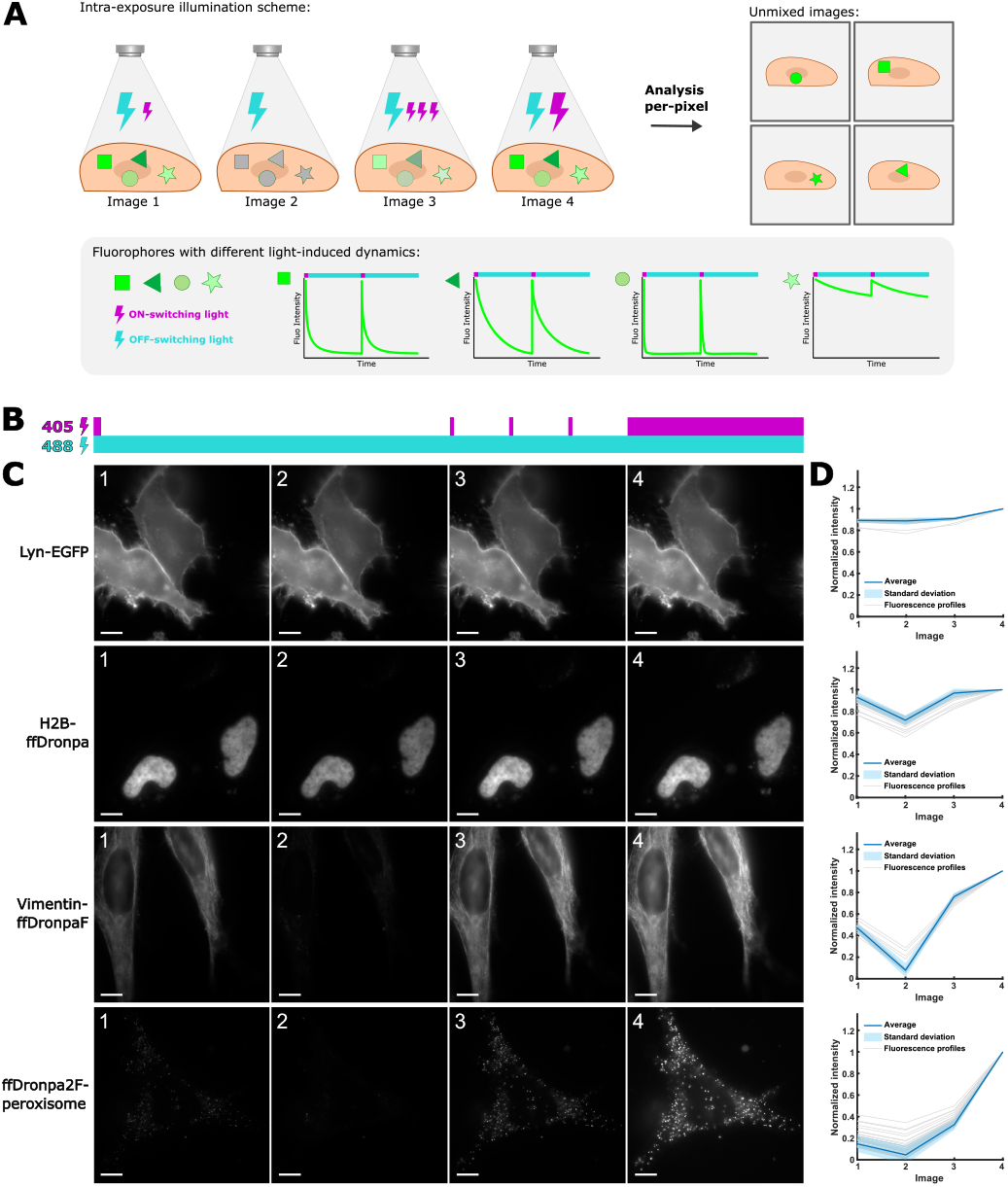
A: Principle of the exNEEMO method. B: Illumination patterns used to probe the distinct photoswitching efficiencies and C: resulting fluorescence images for representative fixed HeLa cells that expressed a single fluorophore. D: Fluorescence profiles (normalized to the fourth image) for all cells expressing single fluorophores. Each line corresponds to a single distinct field-of-view (FOV). Scale bars 10 μm.

In our previous study, the acquisition of two to four fluorescence images with different excitation light modulations were sufficient to classify four different types of fluorophores according to their photoswitching efficiencies. To test this approach in complex mixtures, we expressed the same fluorophores in HeLa cells using different targeting motifs for each label. In particular, we targeted EGFP to the plasma membrane using the Lyn targeting motif (Lyn-EGFP), ffDronpa to the nucleus via fusion with histone-H2B (H2B-ffDronpa), ffDronpaF to vimentin-based fibers via fusion to vimentin (vimentin-ffDronpaF) and ffDronpa2F to peroxisomes (perixosome-ffDronpa2F). To separate such mixtures, the number of acquired images must at least be equal to the number of components in the mixture, and hence we exposed each of these fluorophores to four different illumination modulations (see Figure 1B). The resulting acquisitions revealed both the expected cellular localizations of the labels as well as the pronounced differences in their responses to the illumination (see Figure 1C and Supplementary Movies S1-4)). We then repeated this procedure over many cells expressing single constructs, allowing the measurement of the fluorescence reference profiles for the expressed fluorophore (see Figure 1D). These profiles encode the characteristic response of each type of fluorophore to the illumination modulation.

The characteristic fluorophore responses were apparent also when imaging HeLa cells expressing all four labels simultaneously (see Figure 2A and Supplementary Movie S5), though it is difficult to distinguish the four rsFPs in the measured images by eye due to their strong spatial overlap within the cell. We instead analyzed these images using non-negative least squares [15]. In this approach, the measured image (*X*) is decomposed into the product of the local abundances of each fluorophore (*A*) and the known responses of these fluorophores to the illumination modulation obtained via the single-label experiments (*S*) under a non-negativity constraint [16, 17]. Because the measurement is susceptible to noise, we must also include an error term *E* such that

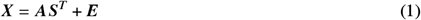

**Figure 2.**
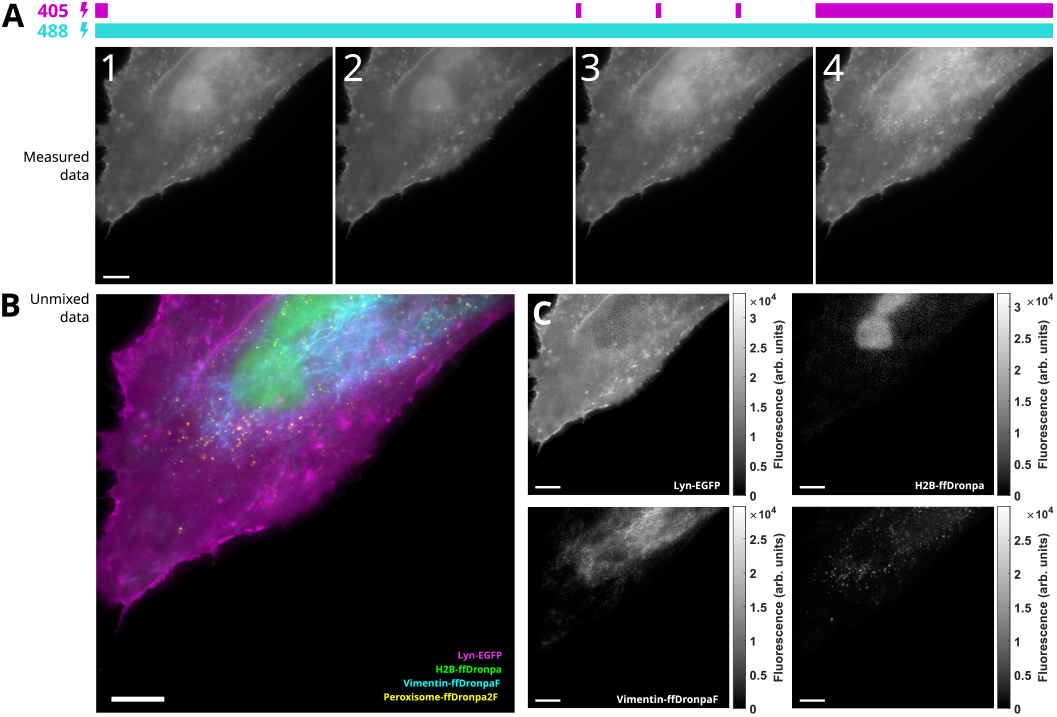
A: HeLa cells expressing all four rsFPs imaged under the specific illumination scheme of violet and cyan light, resulting in a four-image sequence. B: False-color image showing the separation of the different emitter types using the exNEEMO analysis on the raw data shown in panel A. C: Distributions of the individual fluorophores for the same image as in panel B. Scale bars 10 μm.

Given *X* and *S*, the least-squares analysis sets out to solve this system of equations in order to determine *A*. In our experiments, *X, A*, and *E* are of dimensions *N*×4, where *N* is the total number of non-background pixels in a fluorescence image and 4 is the number of acquired images. To fill *X*, the four images acquired with different illumination patterns are simply unfolded by concatenating all of the pixel values from a given image into the individual columns of *X*. The matrix *S* is a 4×4 matrix containing the responses of the fluorophores as determined from the single-fluorophore control measurements. The dimensions of this matrix arise because we have four different fluorophores and four different modulation patterns (fluorescence images). More information on the analysis is provided in the methods section of this work.

Figure 2B shows an example result from this analysis, showing that the contributions of the different fluorophores can be selectively extracted to reveal their expected intracellular targeting. Figure 2C displays each of the calculated fluorophore distributions in separate image plots, making it possible to observe whether there is crosstalk between the different channels. The different fluorophore distributions are readily distinguishable, though some crosstalk is observed between the EGFP and ffDronpa signals. This crosstalk reflects the closer correspondence between the responses of these fluorophores to the illumination modulations (Figure 1D). Supplementary Figure S1 shows additional results of the exNEEMO procedure obtained on randomly-selected cells expressing these constructs.

The exNEEMO analysis decomposes the acquired fluorescence images using the provided single-fluorophore reference profiles, by determining the weighted sum of the reference profiles that best describes the experimental data acquired in each pixel. Scaling the values of a reference profile by a factor *A* therefore results in a rescaling of the calculated fluorophore abundance by a factor 1/*A*. However, regardless of any such scaling, the obtained fluorophore abundances will be proportional to the local fluorophore concentrations, which is exactly the assumption that is applied in the vast majority of fluorescence imaging experiments. The results of our method are therefore compatible with a variety of downstream analysis pipelines and approaches. However, because it relies on the absolute quantification of the emitted fluorescence, the method is sensitive to the presence of background emission and care should be taken to correct for any non-negligible background emission.

A key goal of this work was to evaluate the extent of any uncertainty in the analysis. Figure 3 visualizes the results of our analysis for cells expressing just one type of fluorophore. A visual inspection of example images indeed underscores a high degree of specificity within each of the detected channels (see Figure 3A).

**Figure 3.**
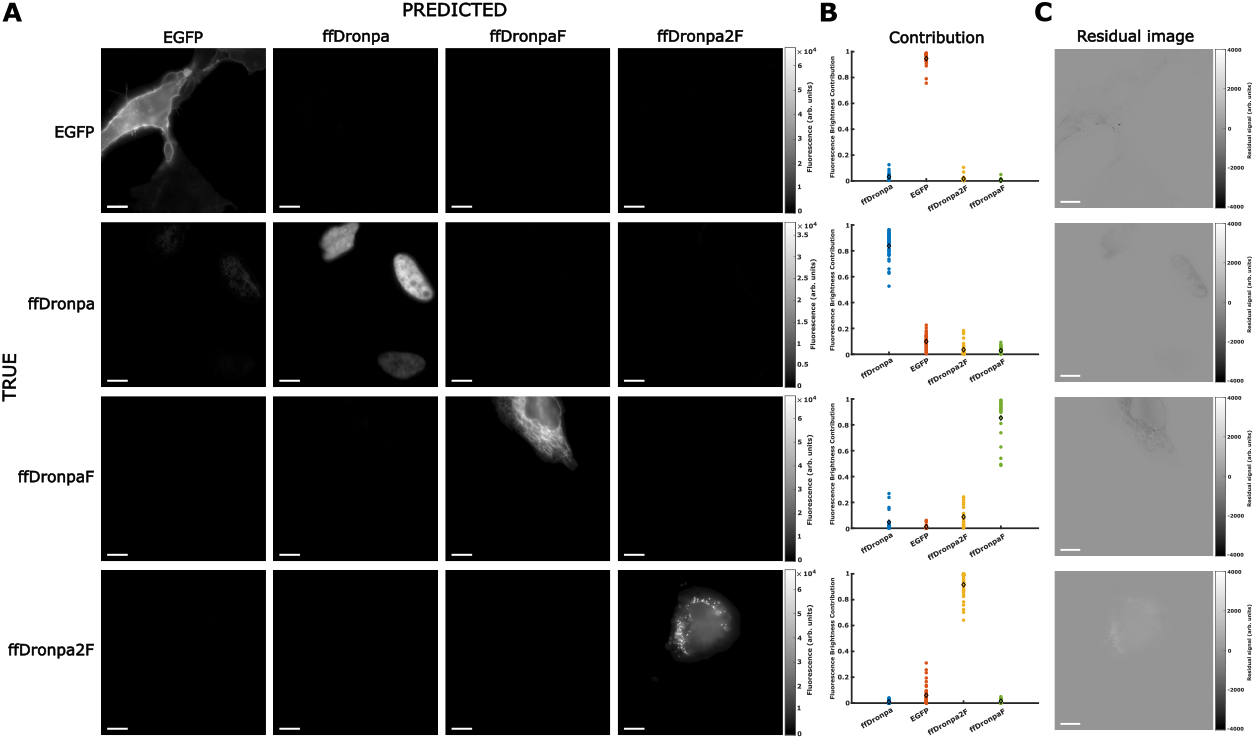
A: exNEEMO analysis of the cells expressing a single rsFP. B: Unmixing accuracies of each rsFP shown as fluorescence brightness contribution plots. One dot represents one FOV. The diamond represents the average. C: Images showing the residual signal when subtracting the fourth image produced by the exNEEMO from the fourth measured fluorescence image. Contributions of all four components were used in the calculation of the residual images.

To obtain statistically-relevant results, we performed an analysis on cells expressing single fluorophores (71 cells distributed over 31 field-of-views (FOVs) for Lyn-EGFP, 83 cells distributed over 39 FOVs for H2B-ffDronpa, 41 cells distributed over 22 FOVs for vimentin-ffDronpaF and 84 cells distributed over 40 FOVs for peroxisome-ffDronpa2F), by calculating the fluorescence brightness contribution determined for each fluorophore for the fourth (brightest) illumination pattern (see Equation 3 and Figure 3B). These showed that on average about 80% to 95% of the signal was correctly assigned by the analysis.

In order to estimate to what extent signals might be unaccounted for or spuriously generated by our analysis, we calculated a residual image for the fourth fluorescence image, showing the difference between the calculated and measured fluorescence intensities (this is equivalent to the refolded matrix *E* – see Equation 1 and the Methods section). Figure 3C visualizes these for the single-fluorophore data, showing that the magnitude of these residuals is within 10% of the calculated fluorescence brightnesses, though a low amount of imperfectly captured structuring is present, particularly for ffDronpaF.

We sought to understand whether this imperfect separation could arise from the variability in the fluorophore response profiles observed between cells expressing the same construct, as was observed in Figure 1D. To do so, we calculated fluorescence brightness contribution plots such as those in Figure 3B but each time varying the response profile of one fluorophore while keeping the remaining three response profiles constant. In other words, Equation 3 was applied repeatedly using one single rsFP response profile from an individual FOV and averaged response profiles for the other rsFPs considering all their respective FOVs. Figure 4 shows an example for varying ffDronpa2F reference profiles, while Supplementary Figure S2 shows this data also when varying the reference profiles of the other fluorophores. We find that the choice of reference profile has an effect on the extracted contributions, but the resulting variation is fairly small, apart from the contribution of specific FOVs that show clearly differing reference profiles (Supplementary Figure S3). These deviating profiles were mainly associated with cells that were dimly fluorescent (see Supplementary Figure S4), that are intrinsically more difficult to analyze, and may also reflect variations induced by the fixation process [18]. These effects can be mitigated by performing a careful validation of the regions or cells selected as part of the single-fluorophore reference data. In practical applications of this method, we suggest the use of residual images such shown in Figure 3C to identify situations where the measured signal cannot be adequately decomposed into the used reference profiles.

**Figure 4.**
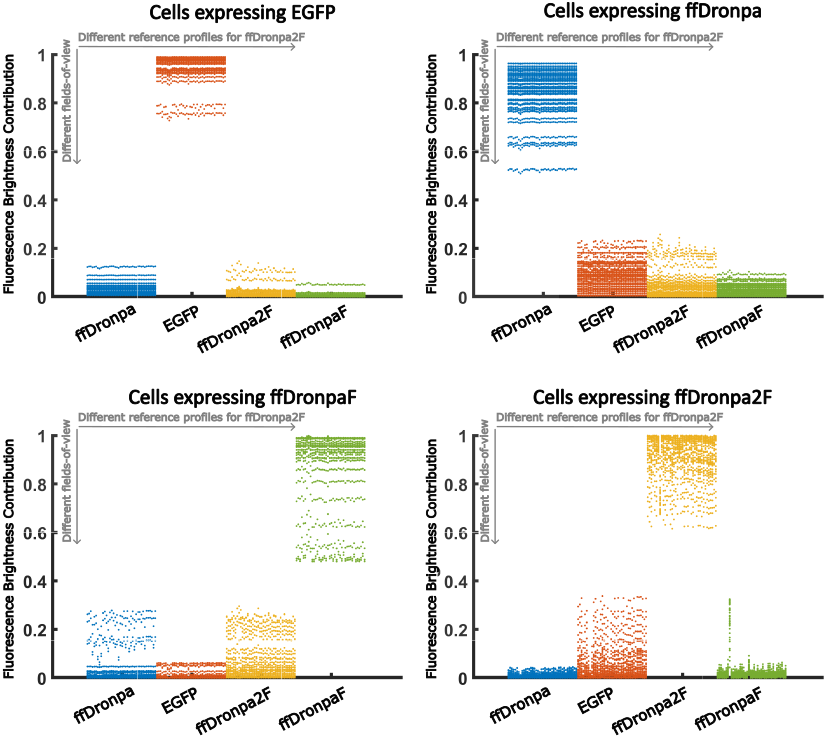
Influence of the reference profile on the fluorophore contribution in the exNEEMO output. Example of a contribution plot showing fluorescence brightness contributions of each rsFP in the single-fluorophore datasets, where reference profiles of ffDronpa2F are varied. Data points along the *y*-axis denote different FOVs.

## Conclusions

In this work, we have developed exNEEMO, an approach for the spatially-resolved separation of fluorophore mixtures based on their light-induced fluorescence dynamics. The core idea is to illuminate the sample with temporally-varying light intensities, to which the fluorophores respond with characteristic emission profiles determined by the nature of the illumination pattern and the kinetics of their fluorescence dynamics. Compared to other approaches, our method has the advantage that it can make use of fluorescence dynamics that may occur too quickly to be resolved directly, but that are well within the rates at which even inexpensive light sources can be modulated. The exNEEMO experiments can in principle be carried out on any microscope that allows for the fast modulation of the excitation light.

Compared to our previous work, which was focused on the classification of samples expressing a single fluorophore each, the methodology in this work can resolve fluorophore mixtures down to the ‘per-pixel’ limit in which the full spatial resolution of the imaging is retained. To do so, we assume that the characteristic fluorophore response profiles are known in advance or can be obtained through measurements of samples expressing a single fluorophore. We analyzed the experimental data using non-negative least squares, which decomposes the measured signal into a weighted sum of the individual fluorophore reference profiles. This analysis does not take into account that the local fluorophore environment or illumination conditions may cause deviations of the reference profiles, which can lead to artifacts in the analysis. Fully addressing such issues will require the development of analysis algorithms that can dynamically adjust the reference profiles to account for changes in probe behavior that may arise within particular (cellular) environments. Such algorithms may well be guided by minimizing the structuring observed in the residual images in order to deliver an emitter separation that accounts for the full measured signal.

In conclusion, using exNEEMO we were able to distinguish four green rsFPs in eukaryotic cells in a per-pixel manner that maintains the full spatial resolution of the imaging. Technically, the principle of exNEEMO can be transferred to photochromic probes of other colors, which would only require wavelength adjustment of the illumination sequence. Similarly to our previous work, the key advantage of the method is the ability to operate on a wide range of fluorescence dynamics since it is not limited by the temporal resolution of the detector. By providing increased multiplexing opportunities using a conceptually simple approach, our method opens new dimensions for high-content imaging in complex (biological) systems.

## Acknowledgements

This work was supported by the Research Foundation-Flanders (FWO) via grants G010723N and G090819N, FLAG-ERA via grant JTC2019 Sensei, and the University of Lille via their International Chair program.

## Methods

### Plasmids

As in our previous work [8], the fluorophores used in this project were EGFP, ffDronpa, ffDronpaF (ffDronpa K45I/F173V), and ffDronpa2F (ffDronpa K45I/M159T/F173V). Plasmids coding for expression of these fluorophores in mammalian cells, *i*.*e*., pcDNA3-Lyn11-EGFP (with N-terminal MGCIKSKGKDSAGGGS targeting sequence), pH2B-ffDronpa (targeted to the histone H2B in the nucleus), pVimentin-ffDronpaF (targeted to the vimentin in the cell cytoskeleton), were already available in the lab. A plasmid for ffDronpa2F targeted to the peroxisome was generated by replacing EGFP with ffDronpa2F with C-terminal SKL sequence in the pEGFP-N1 vector.

### Mammalian cell culture, transfection and fixation

HeLa cells were grown at 37°C in 5% CO_2_ atmosphere in DMEM (ThermoFisher) with GlutaMAX-1 complemented with 10% fetal bovine serum (FBS) and GlutaMAX (ThermoFisher). Cells were seeded in 35-mm glass-bottom dishes (MatTek) and after 24h they were transiently transfected using X-tremeGENE HP DNA Transfection Reagent (Roche) according to the manufacturer’s protocol. 24h after the transfection, cells were washed with 1x Dulbecco’s phosphate-buffered saline (DPBS), pre-fixed with 4% paraformaldehyde (PFA, Electron Microscopy Sciences) solution for 2 min at RT, washed with 1x DPBS and then the fixation was completed with 4% PFA for 12 min at RT. The prefixation step helps to preserve the native protein structures in the cells. Afterwards, cells were incubated with 10% glycine for 10 min at RT and then washed four times with 1x DPBS. Dishes with fixed cells were stored in 4°C in 1x DPBS in the dark until the imaging experiment.

### Widefield image acquisition

Imaging of the fixed cells was performed on a Nikon Eclipse Ti-2 Inverted Microscope (Minato City, Japan) equipped with a 1.4 NA oil immersion objective (100× CFI Apochromat Total Internal Reflection Fluorescence) and a ZT405/488/561/640rpcv2 dichroic mirror with a ZET405/488/561/640 nm emission filter (both Chroma Technology, Bellows Falls, Vermont) in epi-illumination. Two separate lasers at 405 and 488 nm (Oxxius, Lannion, France) were used for excitation. Images were acquired on a PCO.edge 4.2 camera (PCO, Kelheim, Germany) with an exposure time of 50 ms and an optical pixel size of 78.8 nm, using varying pulses of the 405 nm light (∼11 mW transmitted by the objective) in the presence of 488 nm light (∼10mW transmitted by the objective). The excitation light was controlled using an Arduino-compatible microcontroller (Velleman, Gavere, Belgium). Due to the rolling-shutter readout process, the 50 ms camera exposure resulted in a maximal per-image illumination duration of 40 ms since light was applied only when all of the camera pixels were sensitive. To increase the total number of images, the acquisition was performed at equidistant positions over a whole dish using an automated stage loop. FOVs that were out of focus, contained emission that saturated the camera, did not contain any cells or contained clearly degraded cells were excluded from the image analysis.

### Image analysis

The image analysis was carried out by means of non-negative least squares. As was described in the text above (see also [16, 17]), we solve under a non-negativity constraint

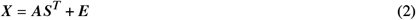

for the matrix *A* that denotes the amount of each fluorophore in every camera pixel, given the measured image *X*, the matrix *S* that contains the reference profiles and *E* an error or residual matrix that describes to what extent the calculated solution deviates from the measured values. In our experiments, *X, A*, and *E* are of dimensions *N*×4, where *N* is the total number of pixels in a fluorescence image and 4 is the number of images. To fill *X*, the four images acquired with different illumination patterns are simply unfolded by concatenating all of the pixel values from a given image into a column of *X*. The matrix *S* is a 4×4 matrix containing the responses of the fluorophores as determined from the single-fluorophore control measurements. The dimensions of this matrix arise because we have four different fluorophores and four different illumination patterns. The actual algorithm used to solve this equation is described in [15].

To reduce the influence of background areas on the analysis, we segmented the acquired images by calculating an average image of all four acquired images, sorting the pixel values, and discarding the pixels that showed the 5% lowest fluorescence brightnesses. To correct for background, the average intensity of these disregarded pixels was then subtracted from the acquired images. The same procedure was performed on data expressing just a single fluorophore. The single-fluorophore reference profiles can then be extracted for each FOV by calculating the average intensity seen in each measured image for the pixels above the threshold. Average reference profiles are determined by averaging over all acquired FOVs for the respective fluorophore.

The contribution graphs in Figure 3B and elsewhere show the averaged fractional contribution of each fluorophore calculated for the single-fluorophore reference data. To calculate the fractional contribution *c*_*k*_ of the *k*-th fluorophore to the fourth (brightest) measured frame, we make use of

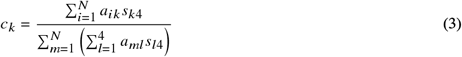

where the indices *l* and *k* refer to the fluorophores ffDronpa, EGFP, ffDronpa2F and ffDronpaF. The summation indices *i* and *m* run over a *N* image pixels included in the analysis. The contribution *c*_*k*_ is calculated independently for each FOV, leading to the individual datapoints shown in Figure 3B. For the contribution graphs with varied reference profiles, we changed the reference profile for one fluorophore and used the average profile for the other three flurophores. The varied profiles were extracted from the individual single-fluorophore FOVs.

## Supporting Information Available

Content of the Supporting Information file:

- Four Supplementary Figures related to image analysis
- Five Supplementary Videos showing measured data

## References

[1] Junyoung Seo, Yeonbo Sim, Jeewon Kim, Hyunwoo Kim, In Cho, Hoyeon Nam, Young-Gyu Yoon, and Jae-Byum Chang. PICASSO allows ultra-multiplexed fluorescence imaging of spatially overlapping proteins without reference spectra measurements. Nature Communications, 13(1):2475, May 2022.

[2] Ilaria Testa, Andreas Schönle, Claas v. Middendorff, Claudia Geisler, Rebecca Medda, Christian A. Wurm, Andre C. Stiel, Stefan Jakobs, Mariano Bossi, Christian Eggeling, Stefan W. Hell, and Alexander Egner. Nanoscale separation of molecular species based on their rotational mobility. Opt. Express, 16(25):21093–21104, Dec 2008. doi: 10.1364/OE.16.021093. URL http://opg.optica.org/oe/abstract.cfm?URI=oe-16-25-21093.

[3] Thomas Niehörster, Anna Löschberger, Ingo Gregor, Benedikt Krämer, Hans-Jürgen Rahn, Matthias Patting, Felix Koberling, Jörg Enderlein, and Markus Sauer. Multi-target spectrally resolved fluorescence lifetime imaging microscopy. Nature Methods, 13(3):257–262, Mar 2016. ISSN 1548-7105. doi: 10.1038/nmeth.3740. URL https://doi.org/10.1038/nmeth.3740.

[4] Gerard Marriott, Shu Mao, Tomoyo Sakata, Jing Ran, David K. Jackson, Chutima Petchprayoon, Timothy J. Gomez, Erica Warp, Orapim Tulyathan, Holly L. Aaron, Ehud Y. Isacoff, and Yuling Yan. Optical lock-in detection imaging microscopy for contrast-enhanced imaging in living cells. Proceedings of the National Academy of Sciences, 105(46):17789–17794, 2008. ISSN 0027-8424. doi: 10.1073/pnas.0808882105. URL https://www.pnas.org/content/105/46/17789.

[5] Jérôme Querard, Tal-Zvi Markus, Marie-Aude Plamont, Carole Gauron, Pengcheng Wang, Agathe Espagne, Michel Volovitch, Sophie Vriz, Vincent Croquette, Arnaud Gautier, Thomas Le Saux, and Ludovic Jullien. Photoswitching Kinetics and Phase-Sensitive Detection Add Discriminative Dimensions for Selective Fluorescence Imaging. Angewandte Chemie International Edition, 54(9):2633–2637, 2015. doi: https://doi.org/10.1002/anie.201408985. URL https://onlinelibrary.wiley.com/doi/abs/10.1002/anie.201408985.

[6] S. Duwé, W. Vandenberg, and P. Dedecker. Live-cell monochromatic dual-label sub-diffraction microscopy by mt-pcSOFI. Chem. Commun., 53:7242–7245, 2017. doi: 10.1039/C7CC02344H. URL http://dx.doi.org/10.1039/C7CC02344H.

[7] Daniel Wüstner. Dynamic Mode Decomposition of Fluorescence Loss in Photobleaching Microscopy Data for Model-Free Analysis of Protein Transport and Aggregation in Living Cells, 2022. ISSN 1424-8220. URL https://doi.org/10.3390/s22134731.

[8] Hana Valenta, Siewert Hugelier, Sam Duwé, Giulia Lo Gerfo, Marcel Müller, Peter Dedecker, and Wim Vandenberg. Separation of spectrally overlapping fluorophores using intra-exposure excitation modulation. Biophysical Reports, 1(2):100026, Dec 2021. ISSN 2667-0747. URL https://www.sciencedirect.com/science/article/pii/S2667074721000264.

[9] Raja Chouket, Agnès Pellissier-Tanon, Aliénor Lahlou, Ruikang Zhang, Diana Kim, Marie-Aude Plamont, Mingshu Zhang, Xi Zhang, Pingyong Xu, Nicolas Desprat, Dominique Bourgeois, Agathe Espagne, Annie Lemarchand, Thomas Le Saux, and Ludovic Jullien. Extra kinetic dimensions for label discrimination. Nature Communications, 13(1):1482, Mar 2022. ISSN 2041-1723. doi: 10.1038/s41467-022-29172-0. URL https://doi.org/10.1038/s41467-022-29172-0.

[10] Yuling Yan, M Emma Marriott, Chutima Petchprayoon, and Gerard Marriott. Optical switch probes and optical lock-in detection (OLID) imaging microscopy: high-contrast fluorescence imaging within living systems. Biochem J, 433(3):411–22, Feb 2011. doi: 10.1042/BJ20100992.

[11] Jung-Cheng Hsiang, Amy E Jablonski, and Robert M Dickson. Optically modulated fluorescence bioimaging: visualizing obscured fluorophores in high background. Acc Chem Res, 47(5):1545–54, May 2014. doi: 10.1021/ar400325y.

[12] Gerardo Abbandonato, Barbara Storti, Giovanni Signore, Fabio Beltram, and Ranieri Bizzarri. Quantitative optical lock-in detection for quantitative imaging of switchable and non-switchable components. Microsc Res Tech, 79 (10):929–937, Oct 2016. doi: 10.1002/jemt.22724.

[13] Jérôme Quérard, Ruikang Zhang, Zsolt Kelemen, Marie-Aude Plamont, Xiaojiang Xie, Raja Chouket, Insa Roemgens, Yulia Korepina, Samantha Albright, Eliane Ipendey, Michel Volovitch, Hanna L. Sladitschek, Pierre Neveu, Lionel Gissot, Arnaud Gautier, Jean-Denis Faure, Vincent Croquette, Thomas Le Saux, and Ludovic Jullien. Resonant out-of-phase fluorescence microscopy and remote imaging overcome spectral limitations. Nature Communications, 8(1):969, 2017.

[14] Agnès Pellissier-Tanon, Raja Chouket, Ruikang Zhang, Aliénor Lahlou, Agathe Espagne, Annie Lemarchand, Vincent Croquette, Ludovic Jullien, and Thomas Le Saux. Resonances at Fundamental and Harmonic Frequencies for Selective Imaging of Sine-Wave Illuminated Reversibly Photoactivatable Labels. ChemPhysChem, 23 (23):e202200295, 2022. doi: https://doi.org/10.1002/cphc.202200295. URL https://chemistry-europe.onlinelibrary.wiley.com/doi/abs/10.1002/cphc.202200295.

[15] Rasmus Bro and Sijmen De Jong. A fast non-negativity-constrained least squares algorithm. Journal of Chemometrics: A Journal of the Chemometrics Society, 11(5):393–401, 1997.

[16] Cyril Ruckebusch, Beata Walczak, and Lutgarde Buydens. Resolving spectral mixtures: With applications from ultrafast time-resolved spectroscopy to super-resolution imaging. Data Handling in Science and Technology. Elsevier, 2016.

[17] Siewert Hugelier, Raffaele Vitale, and Cyril Ruckebusch. 4.17 - Image Processing in Chemometrics. In Steven Brown, Romà Tauler, and Beata Walczak, editors, Comprehensive Chemometrics (Second Edition), pages 411–436. Elsevier, Oxford, second edition edition, 2020. ISBN 978-0-444-64166-3. doi: https://doi.org/10.1016/B978-0-12-409547-2.14597-4. URL https://www.sciencedirect.com/science/article/pii/B9780124095472145974.

[18] Katharina N Richter, Natalia H Revelo, Katharina J Seitz, Martin S Helm, Deblina Sarkar, Rebecca S Saleeb, Elisa D’Este, Jessica Eberle, Eva Wagner, Christian Vogl, Diana F Lazaro, Frank Richter, Javier Coy-Vergara, Giovanna Coceano, Edward S Boyden, Rory R Duncan, Stefan W Hell, Marcel A Lauterbach, Stephan E Lehnart, Tobias Moser, Tiago F Outeiro, Peter Rehling, Blanche Schwappach, Ilaria Testa, Bolek Zapiec, and Silvio O Rizzoli. Glyoxal as an alternative fixative to formaldehyde in immunostaining and super-resolution microscopy. The EMBO Journal, 37(1):139–159, 2018. doi: https://doi.org/10.15252/embj.201695709.

