## Supplementary material for "Per-pixel unmixing of spectrally overlapping fluorophores using intra-exposure excitation modulation": SI

July 21, 2023

### Supplementary Figures

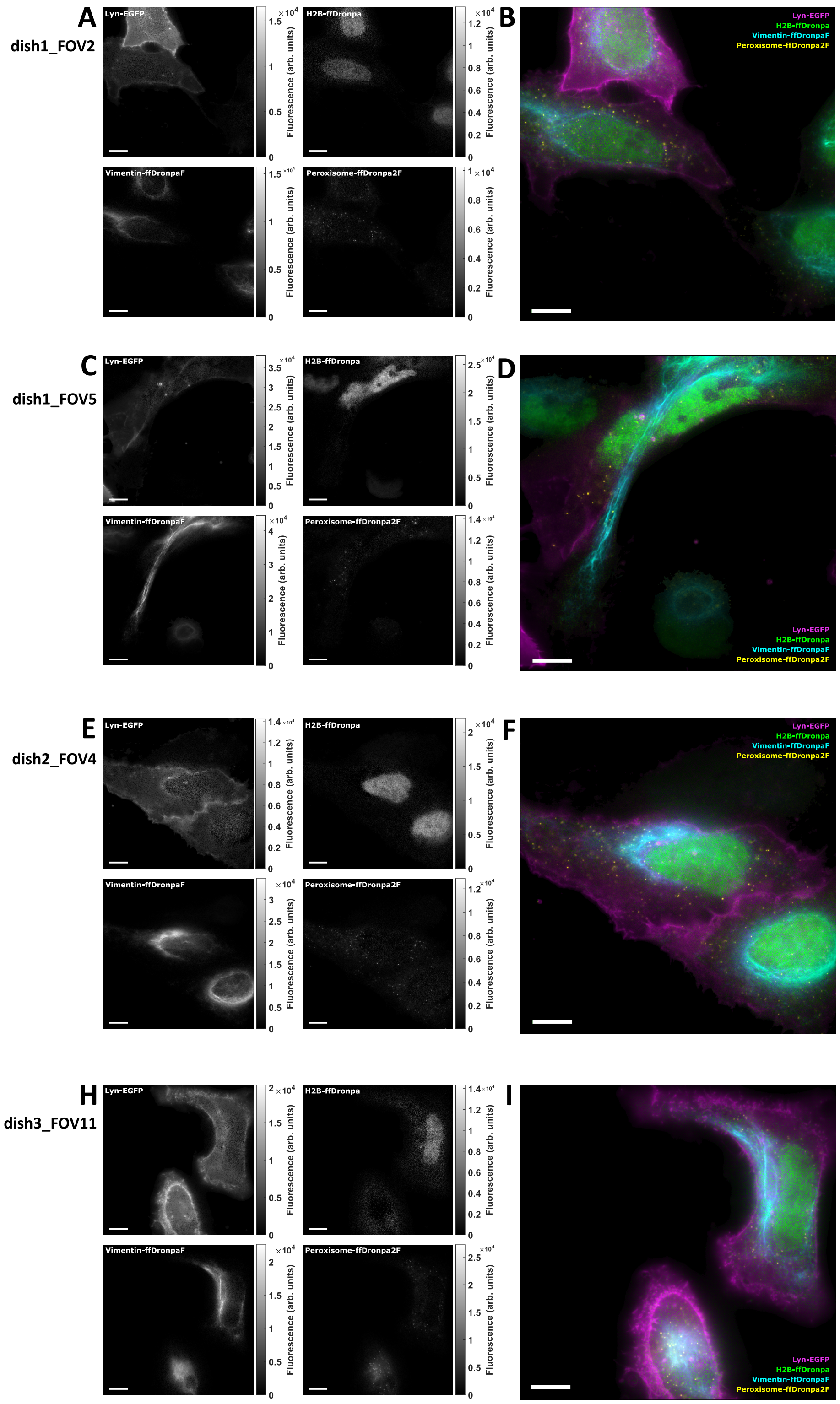


Figure S1: Random selection of FOVs of HeLa cells expressing four rsFPs. A, C, E, H: exNEEMO output - unmixed rsFPs (EGFP, ffDronpa, ffDronpaF and ffDronpa2F) revealing four different cell structures, *i.e.*, plasma membrane, nucleus, vimentin and peroxisomes, respectively. B, D, F, I: Merge of false color images produced by exNEEMO. Scale bars 10 µm.

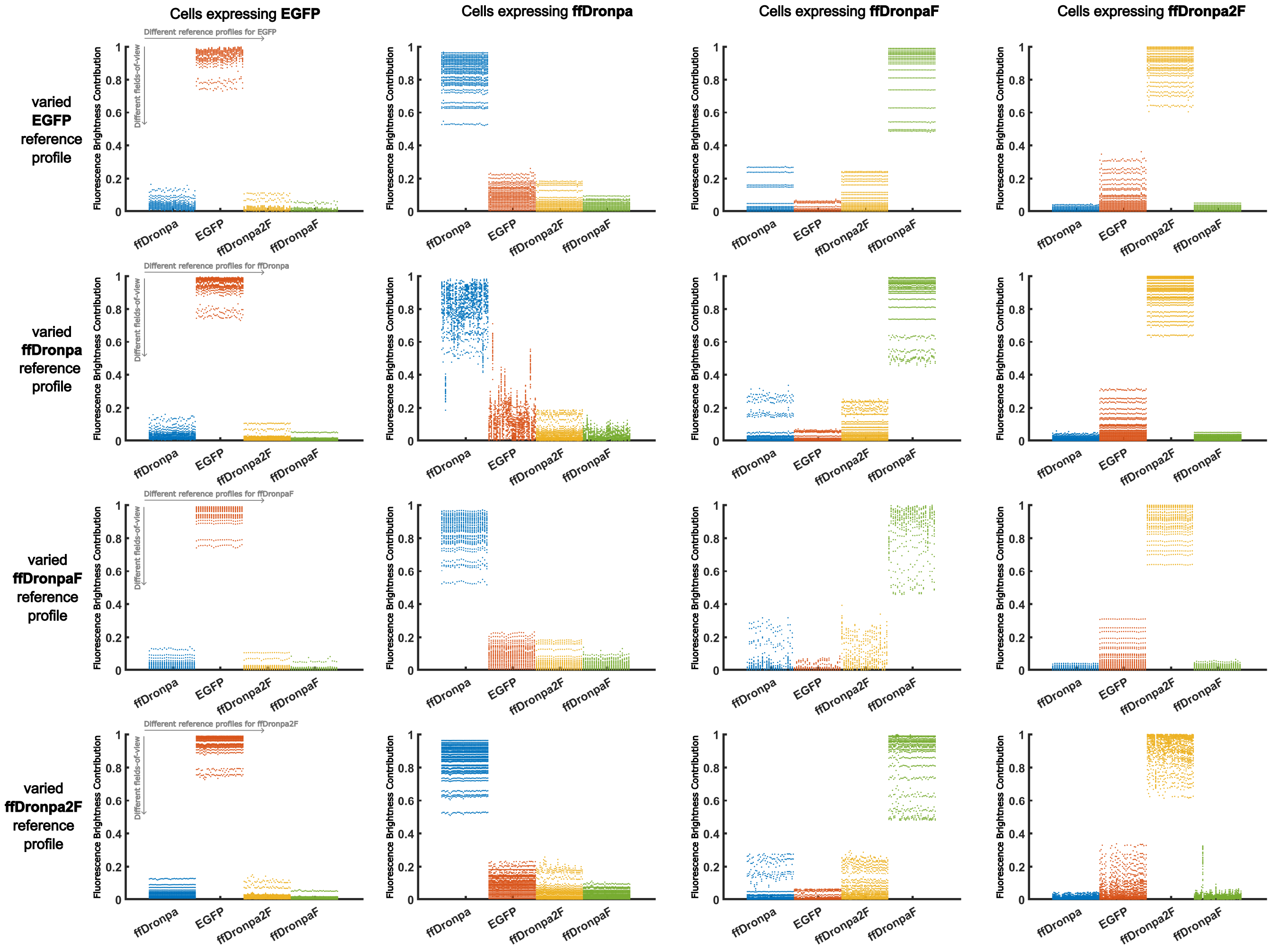


Figure S2: Plots showing the calculated fluorescence brightness contributions for each rsFP in cells expressing just a single fluorophore. In each contribution plot, the reference profile of one rsFP is varied (along the $x$-axis) while using the average reference profiles for the other three rsFPs. The varied profile is extracted from only one single-fluorophore FOV. Depending on the fluorophore, 22 to 40 profile variations were considered. Every data point denotes one FOV (along the $y$-axis) for which the fluorescence brightness contribution was determined.

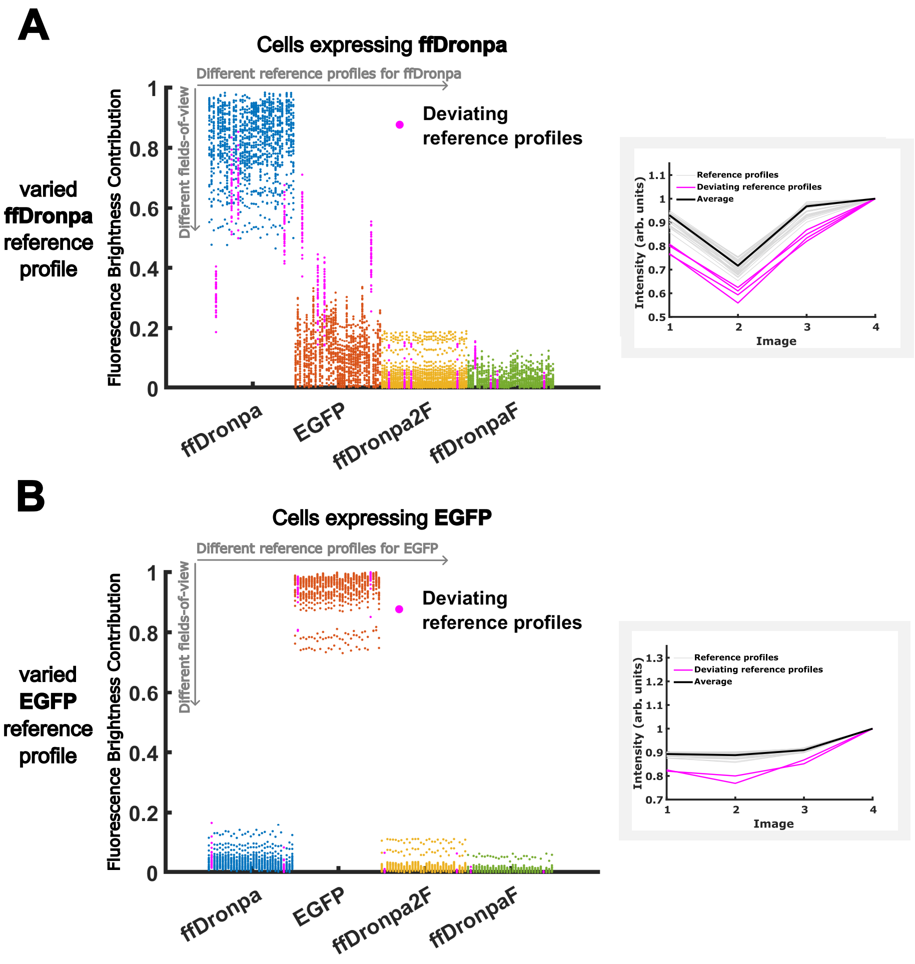


Figure S3: Detailed analysis of contribution plots of ffDronpa (A) and EGFP (B) (from Figure S2). Data points highlighted in magenta show unmixing results when using the magenta-colored (deviating) reference profiles shown in the graphs on the right.

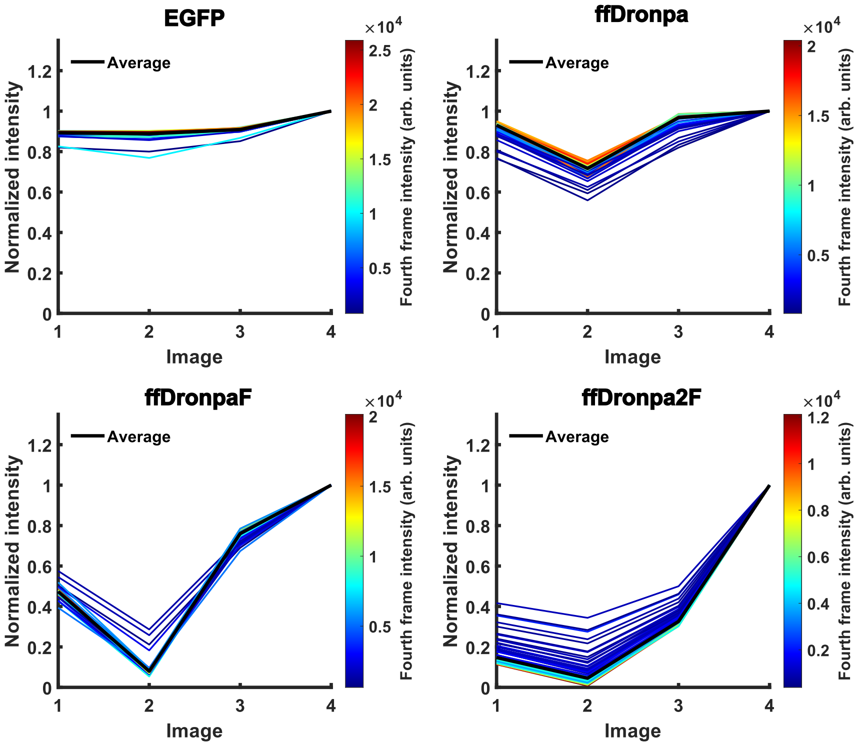


Figure S4: Fluorescence profiles (normalized to the fourth image) for all cells expressing only a single type of fluorophore. Each line shows a single distinct field-of-view, colorcoding shows the observed fluorescence brightness.

### Supplementary Videos

The supplementary material of this manuscript includes 5 videos:

- MovieS1_Lyn-EGFP_HeLa_cells : HeLa cells expressing EGFP targeted to the plasma membrane and illuminated with 405 nm and 488 nm light in a four-image sequence shown in Figure 1B. Image numbers are indicated in the movie.

- MovieS2_H2B-ffDronpa_HeLa_cells : HeLa cells expressing ffDronpa targeted to the histone H2B in the nucleus and illuminated with 405 nm and 488 nm light in a four-image sequence shown in Figure 1B. Image numbers are indicated in the movie.

- MovieS3_vimentin-ffDronpaF_HeLa_cells : HeLa cells expressing ffDronpaF targeted to cell cytoskeleton and illuminated with 405 nm and 488 nm light in a four-image sequence shown in Figure 1B. Image numbers are indicated in the movie.

- MovieS4_Prx-ffDronpa2F_HeLa_cells : HeLa cells expressing ffDronpa2F targeted to the peroxisomes and illuminated with 405 nm and 488 nm light in a four-image sequence shown in Figure 1B. Image numbers are indicated in the movie.

- MovieS5_Mix_Hela_dish2_FOV4 : HeLa cells co-expressing EGFP, ffDronpa, ffDronpaF and ffDronpa2F targeted to the cell structures mentioned before and illuminated with 405 nm and 488 nm light in a four-image sequence shown in Figure 1B. Image numbers are indicated in the movie.
